# Gut microbiota of Brazilian *Melipona* stingless bees: dominant members and their localization in different gut regions

**DOI:** 10.1101/2025.06.03.657762

**Authors:** Amanda Tristao Santini, Alan Emanuel Silva Cerqueira, Nancy A. Moran, Helder Canto Resende, Weyder Cristiano Santana, Sergio Oliveira de Paula, Cynthia Canedo da Silva

## Abstract

The gut microbiome of eusocial corbiculate bees, which include honeybees, bumblebees, and stingless bees, consists of anciently associated, host-specific bacteria that are vital for bee health. Two symbionts, *Snodgrassella* and *Gilliamella*, are ubiquitous in honeybees and bumblebees. However, their presence varies in the stingless bee clade (Meliponini), a group with pantropical distribution. They are absent or rare in the diverse genus *Melipona*, indicating a shift in microbiota composition in this lineage. To identify the main members of the *Melipona* microbiota, we combined newly collected and published data from field-collected individuals of several species. Additionally, we identified the localization of the dominant microbiota members within the gut regions of *Melipona quadrifasciata anthidioides*. The dominant microbiota of *Melipona* species includes members of the genera *Bifidobacterium, Lactobacillus, Apilactobacillus, Floricoccus*, and *Bombella*. Among these, *Apilactobacillus* and *Bombella* dominate in the crop, whereas *Apilactobacillus* and other members of the Lactobacillaceae dominate the ventriculus. The ileum lacks *Snodgrassella* or *Gilliamella* but contains a putative new symbiont close to *Floricoccus*, as well as strains of *Bifidobacterium*, Lactobacillaceae (including *Apilactobacillus*), and *Bombella*. The rectum is dominated by *Bifidobacterium* and *Lactobacillus*. In summary, the *Melipona* microbiota is compositionally distinct but shows spatial organization paralleling that of other eusocial corbiculate bees.

## Introduction

The relationship between insects and microorganisms is vital for the diversification and evolutionary success of insects [1]. Social bees host a diverse and specific gut microbiota that includes core members found across multiple bee species, as well as environmental bacteria [2]. These microorganisms play a crucial role in maintaining the health of bees [3,4]. They acquire their microbiome through social interactions with other colony members, exposure to their surroundings, and their diet [2,5,6].

Eusocial corbiculate bees comprise three clades, the honeybees (genus *Apis*), bumblebees (genus *Bombus*), and stingless bees (tribe Meliponini) [7]. Their gut microbiomes contain anciently associated, host-specific bacteria that can contribute to bee health [2,8,9]. In guts of both honeybees and bumblebees, *Snodgrassella* and *Gilliamella* strains dominate in the ileum, while *Bombilactobacillus, Lactobacillus melliventris*, and *Bifidobacterium* strains dominate in the rectum [2,10,11]. In the stingless bees, *Snodgrassella* and *Gilliamella* vary in occurrence, having been lost/rare in some clades, including the large Neotropical genus, *Melipona* [6,9,12–15]. In *Melipona,* the functional roles of *Snodgrassella* and *Gilliamella* have been speculated to be replaced by new symbionts [12], including a member of the family Streptococcaceae, close to *Floricoccus* and consistently found in *Melipona* species [12–14]. Here, we inferred the dominant members of the *Melipona* Illiger, 1806 microbiome by combining newly collected and published data on gut bacterial communities of field-collected individuals of several Brazilian stingless bees’ species. In addition, we determined the localization of the dominant bacteria to different gut regions within the species *Melipona quadrifasciata anthidioides* Lepeletier, 1836. Our results add to the understanding of the shifts in microbiota structure that have occurred in *Melipona*, including a possible replacement of *Snodgrassella* and *Gilliamella* by new symbionts.

### Methodology

The sample collection was authorized by the Brazilian Environment Ministry (SISBIO/ICMBIO authorization number 87892-1). To infer the dominant members of *Melipona* microbiome, we collected bees from ten (10) populations (i.e. bees from the same species living at the same sampling location) across different locations in Brazil. The populations consisted of two *Melipona* species identified by comparison with known specimens and/or taxonomic keys [16] and five morphotypes whose identification was not confirmed (referred to as “*Melipona* cf. *= conferatum*”). The number of colonies collected per population varied based on availability in each location, as shown in Supplementary S1 Table. Each colony consisted of a beekeeping box, from which forager bees were collected from the entrance and placed in sterile tubes containing 95% ethanol. Five (5) bees from each box were dissected using sterile forceps with a stereoscopic microscope, and their guts comprised a pooled sample.

To assess the microbial diversity in each gut region we selected the *M. quadrifasciata* species the most studied *Melipona* species so far [6,12, 13, 15], highly available in our university. We collected forager bees from 3 different colonies in Viçosa – MG, Brazil (Supplementary S1 Table), and dissected the gut of ten bees into four regions: crop, ventriculus, ileum, and rectum. Each region was treated as a separate sample, totalizing 40 samples (one rectum sample was later discarded).

For all samples in this study, the total DNA was extracted using the NucleoSpin soil kit (Macherey-Nagel), preceded by a proteinase K treatment for 2 hours at 56 ºC, as described in previous work [12]. After extraction, the DNA was submitted for 250 bp paired-end amplicon sequencing at Novogene Corporation Inc (Sacramento, CA, USA) using an Illumina NovaSeq 6000 System. The primer pair 341F (CCTAYGGGRBGCASCAG) and 806R (GGACTACNNGGGTATCTAAT) was used to target the 16S rRNA V3-V4 regions. The data were processed together with previously published data (SRA accession #PRJNA678404) [12] using the DADA2 package (version 1.28) [17] in R 4.3.1, following the pipeline available at https://benjjneb.github.io/dada2/tutorial.html. The taxonomy was assigned to ASVs using a trained SILVA database (version 138.1 from November 2020) for bacteria. For data analysis, we used the R package “mctoolsr” version 0.1.1.9 (available at https://github.com/leffj/mctoolsr), “vegan” version 2.6-4 [18], and “ggplot2” version 3.4.2 [19]. The data was rarefied to reduce bias and make it easier to detect meaningful differences in community composition. Furthermore, the most abundant and *core-like* ASVs (ASVs present in all bee populations analyzed) were submitted to BLASTN similarity searches against GenBank at NCBI Reference Sequence Database at which we could identify and download sequences from isolates aligned to them. Downloaded sequences were aligned using MAFFT 7 [20], and the Maximum Likelihood phylogenetic tree was made with a bootstrap of 1000 replications using IQ-TREE 2 [21]. By this approach we could determine the possible origin of dominant ASVs in *Melipona* (S3 Table, S4 Fig.).

## Results

The microbiota of Brazilian *Melipona* bees is more similar within the same subgenera and biome (S1 Fig.), consistently comprising Acetobacteraceae, Bifidobacteriaceae, Lactobacillaceae, and Streptococcaceae (S2 Fig). Genera present in all samples include *Apilactobacillus, Bifidobacterium, Bombella, Commensalibacter, Floricoccus, Lactobacillus*, and *Neokomagataea*. A few samples contain other environmental genera, such as *Prevotella, Rosenbergiella*, and *Weissella* (Fig. 1, S3 Fig.).

**Fig 1.**
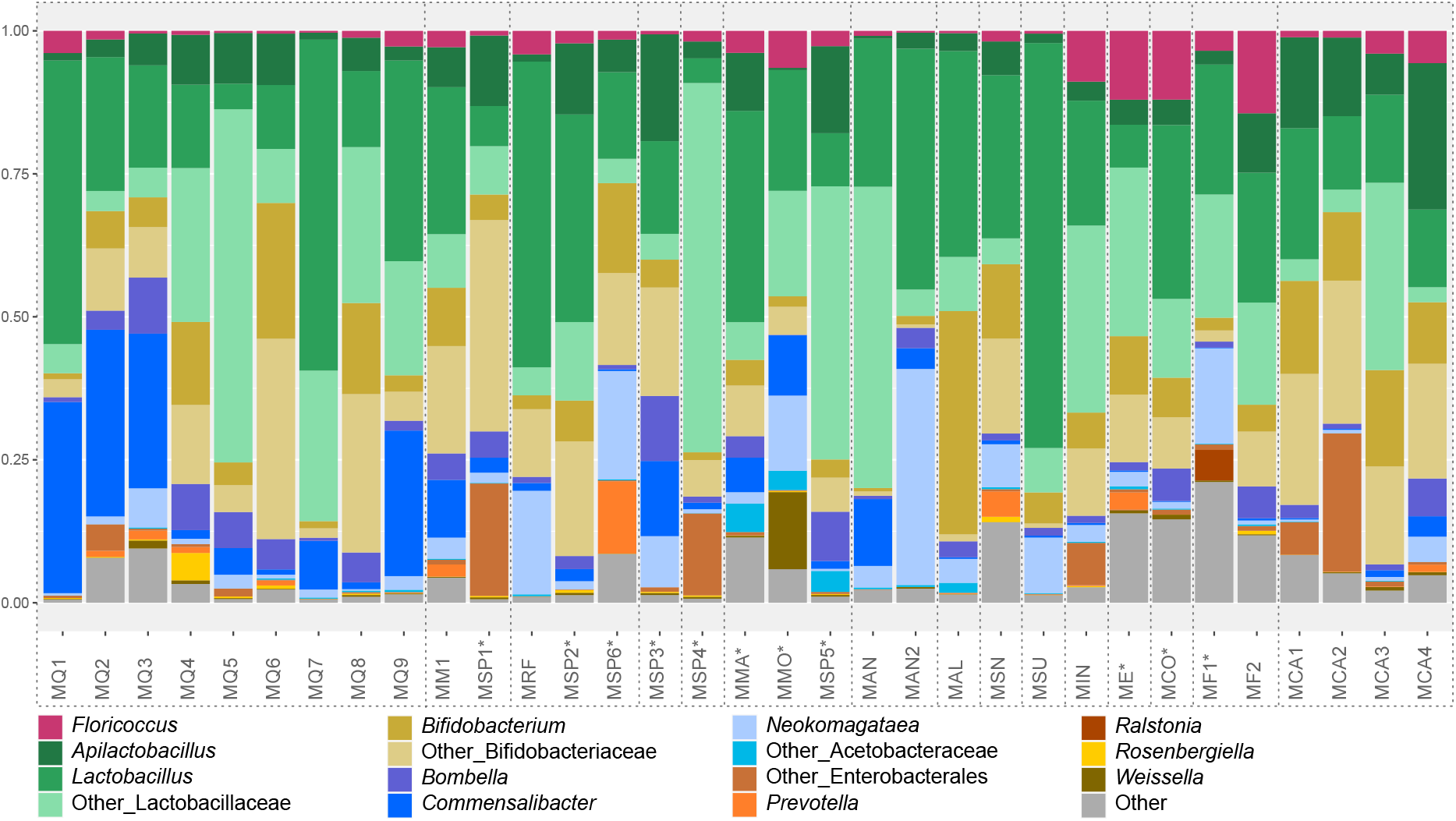
Mean relative abundance of gut bacterial genera in *Melipona* populations classified using SILVA database. Each column represents the mean relative abundance of each population (represented in S2 Fig). ‘Other_Lactobacillaceae’ refers to bacteria assigned to Lactobacillaceae that could not be identified at the genus level. Similarly, ‘Other_Acetobacteraceae’ refers to bacteria assigned to Acetobacteraceae that could not be identified at the genus level. ‘Other_Enterobacterales’ refers to bacteria only identified at the order level. ‘Other’ are bacteria in lower abundance. See Table S1 for population and collection information. Populations grouped by dotted lines are considered from the same *Melipona* species. *Species whose identification was not confirmed.

Concerning the *Melipona quadrifasciata* gut regions, the ileum presents a higher alpha diversity (Shannon index, S4 Fig) compared with the other gut regions. However, there is no statistical difference in the richness index (S4 Fig) among the different gut regions. The NMDS based on the Bray-Curtis dissimilarity matrix separated the samples by region but not by source colony (Fig. 2C), and PERMANOVA analysis revealed significant differences among gut regions, except between ventriculus and ileum (S2 Table).

**Fig 2.**
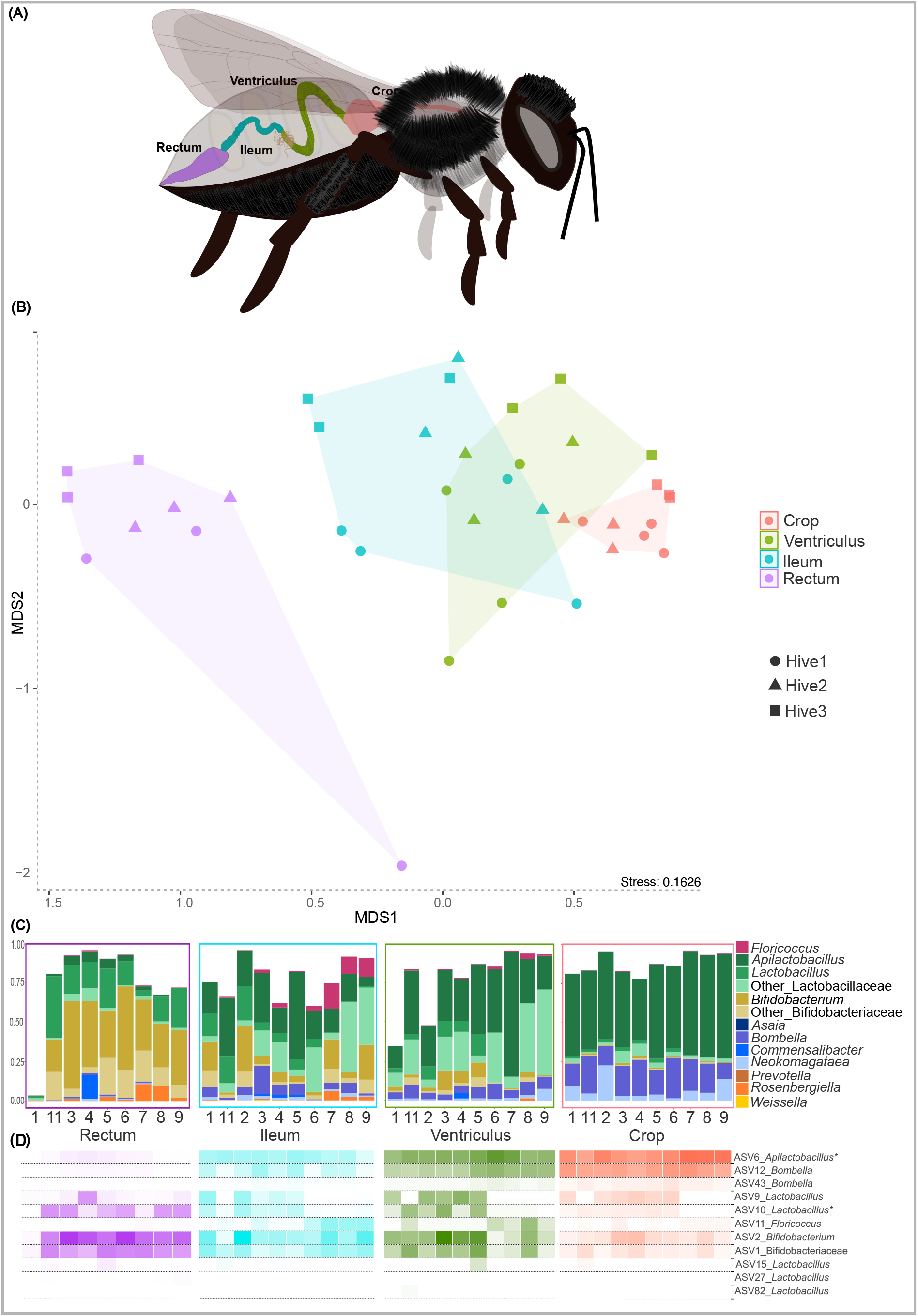
Microbial community of gut regions of *M. quadrifasciata anthidioides*. (A) Schematic figure of *Melipona quadrifasciata* gut. (B) NMDS based on ASV relative abundance (Bray-Curtis dissimilarity) in gut regions of bees from three colonies. (C) Relative abundance of dominant bacterial genera classified using SILVA database, in each gut region. (D) Heatmap of *Melipona core-like* ASVs in each gut region classified using SILVA database. *^1^ASV6 was classified as *Apilactobacillus* using SILVA database but formed a clade with *Nicoliella* using Genbank Nucleotide Database sequences (see S4 Fig). *^2^ASV11 was classified as *Floricoccus* using SILVA database but formed a clade with yet undescribed Streptococacceae isolates close to *Floricoccus* using Genbank Nucleotide Database sequences (see S4 Fig).

The genera that are more abundant in *Melipona* generally compose more than 70% of the community in individual gut regions. However, gut regions have distinct compositions. The crop is dominated by *Apilactobacillus, Bombella*, and *Neokomagataeae* (Fig. 2B); the ventriculus by *Apilactobacillus*, other Lactobacillaceae, *Bombella,* and Bifidobacteriaceae; the ileum by Lactobacillaceae (including *Apilactobacillus* and *Lactobacillus*), Bifidobacteriaceae (including *Bifidobacterium*), *Bombella,* and *Floricoccus*; and the rectum by Bifidobacteriaceae (including *Bifidobacterium*) and Lactobacillaceae (including *Apilactobacillus* and *Lactobacillus*). Interestingly, a sequential decrease is observed for the relative abundance of *Apilactobacillus* from the crop to the rectum. *Bombella* is also more abundant in the crop compared to ventriculus and ileum. Alternatively, an opposite trend is observed for *Bifidobacterium* and other Bifidobacteriaceae, which increase their relative abundance from the ventriculus to the rectum, where they are the main colonizers along with *Lactobacillus*. Of the total 1,690 ASVs in the samples, 11 ASVs are present in all species of *Melipona* and are considered the *core-like* microbiota members (Fig. 2C). These 11 ASVs are related to *Bifidobacterium, Bombella, Floricoccus, Lactobacillus*, and *Apilactobacillus*. We created phylogenies for *Melipona* dominant and most abundant ASVs to differentiate between bacteria consistently associated with bees and bacteria found in other environments (S5 Fig). ASVs of *Lactobacillus, Bombella* and *Bifidobacterium* groups in *Melipona* are related to those found in other bees, including isolates from bumblebees [22]. The *Floricoccus* ASV, although close to environmental isolates, formed a distinct clade together with strains previously isolated from *Melipona* [14]. Similarly, the *Apilactobacillus* ASVs are closely related to *Nicoliella spurrieriana*, a bacterium isolated from *Tetragonula carbonaria*, an Australian stingless bee [23]. These observations point towards two possible stingless bee-associated new clades (Fig. 2C, S5 Fig).

Among the *core-like* ASVs, ASV6 (*Apilactobacillus*) and ASV12 (*Bombella*) are the most prevalent in both crop and ventriculus. ASV9 and AS10 (*Lactobacillus*) are more abundant in the ventriculus, ileum and rectum, while ASV11 (*Floricoccus*) is more prominent in the ileum. Additionally, ASV1 and ASV2 (Bifidobacteriaceae and *Bifidobacterium*, respectively) show increased relative abundance in the ileum and rectum. Although the other core-like ASVs have lower abundances in each gut region, they are consistently present in all analyzed regions of *M. quadrifasciata*.

Overall, Brazilian *Melipona* bees lack core bacterial lineages typically associated with honeybees, including *Gilliamella, Snodgrassella* and *Bombilactobacillus* (former Firm-4). Instead, they have acquired new putative core-like bacterial lineages, such as *Floricoccus* (Fig. 3).

**Fig 3.**
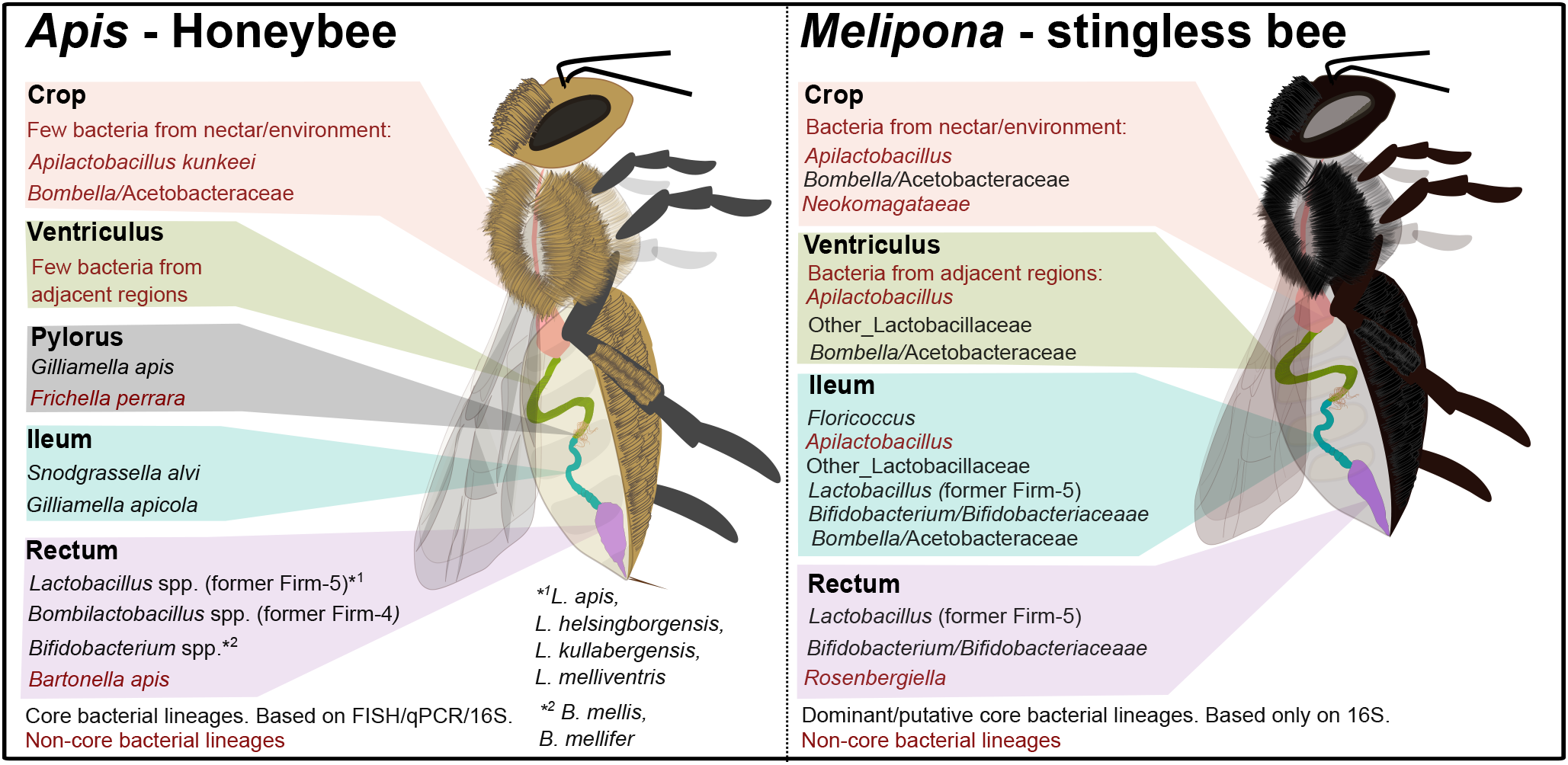
Comparative schematic of gut microbiota composition in Apis and Brazilian Melipona bees across different gut regions.

## Discussion

The microbiota of *Melipona* differs from that of other eusocial bees, with rare/no occurrence of the symbionts *Snodgrassella* and *Gilliamella*, corroborating previous observations [6,12– 14,24]. The Brazilian *Melipona* microbiota is mainly composed of *Bifidobacterium, Lactobacillus, Apilactobacillus, Floricoccus,* and *Bombella*, as they are present in all bee populations analyzed. This study marks the first comprehensive analysis of the *Melipona* gut regions and their microbial composition. We specifically chose to analyze *M. quadrifasciata* due to its widespread occurrence in Brazil, and its role in honey production and agricultural pollination. In addition, the abundance of research available on this species [6,12,13,15] enabled us to assess the consistency between the microbial communities across the gut regions and the dominant members of the *M. quadrifasciata* microbiome. Notably, the primary microbes found in the crop, the sugar-rich honey stomach of bees, are *Apilactobacillus* and *Bombella* [23,25]. The ventriculus also has *Apilactobacillus* and *Bombella* as well as several Lactobacillaceae, including the *Lactobacillus core-like* ASV9 and ASV10. These microorganisms are fructophilic species commonly associated with the hive environment and honey [6,26]. In addition, these findings align with other studies on bee gut microbiota, which have shown that the anterior region of the gut, including the crop and ventriculus, hosts both environmental and transient microbiota [27].

In other social bees, over 90% of the gut microbiota is found in the hindgut, consisting of ileum and rectum [10]. In *M. quadrifasciata*, the rectum is dominated by *Bifidobacterium* and *Lactobacillus*, as observed for the *core* microbiota of other eusocial corbiculate bees [5,28], but the ileum has a very different composition. The *M. quadrifasciata* ileum contains the putative new symbiont close to *Floricoccus* and already isolated from *Melipona* [14] as well as strains of *Bifidobacterium*, Lactobacillaceae (including *Apilactobacillus*), and *Bombella.* In contrast, in honeybees, *Bombella* and *Apilactobacillus* are largely limited to the crop [22,29,30]. Potentially, the distinct ileum community of *Melipona* carries out the same metabolic and defensive functions as the *Snodgrassella/Gilliamella*-dominated ileum community of honeybees and bumblebees. Further experimental studies using microbial isolates and bee colonization assays will be done to explore this issue.

## Acknowledgments

We acknowledge the UFV, and financial support from CNPq, CAPES – Finance Code 001, FAPEMIG (Finance Code APQ – 03029-21) and from US NIH award R35GM131738 to NAM and AESC. We thank Anderson Alexandre, Ricardo Marinho Gomes, and Eduardo da Costa Tavares for providing bees.

## Data Availability

The 16S rRNA gene amplicon sequencing raw data were deposited in the NCBI BioProject database under the accession number PRJNA1076254.

## Supporting information

**S1 File. Supporting tables and figures.** This PDF contains (1) S1 Table. Information of collection, species name and source of the *Melipona* samples analyzed in the present work. (2) S2 Table. PERMANOVA based on the Bray-Curtis dissimilarity matrix comparing the differences in the microbial community composition between the gut regions of *M. quadrifasciata*. (3) S3 Table. GenBank sequences used for analysis. (4) S1 Figure. NMDS plot based on ASV relative abundance using a Bray-Curtis dissimilarity matrix, illustrating bacterial community composition across different *Melipona* species and biomes. Colors represent bee species, with color groupings indicating *Melipona* subgenera: orange – *Melipona*, green – *Michmelia*, blue – *Eomelipona*, and pink – *Melikerria*. Point shapes denote the biome of origin. (5) S2 Figure. Most abundant families in *Melipona* spp. gut microbiota. Each sample represents a pool of 5 bees per box per site of study. ASVs are ordered and colored at the family level, with low abundant ASVs grouped as ‘Other’. (6) S3 Figure. Most abundant genera in *Melipona* spp. gut microbiota. Each sample represents a pool of 5 bees per box per site of study. ASVs are ordered and colored at the genus level, with low abundant ASVs grouped as ‘Other’. (7) S4 Figure. Bacterial alpha diversity of the gut regions of *M. quadrifasciata*. The alpha diversity was expressed using the Shannon and richness indexes. A Kruskal-Wallis test (p < 0.05) was conducted, followed by a post-hoc pairwise Dunn test to compare each gut part, showing only the significant results. (8) S5 Figure. Phylogenetic trees of the most abundant ASVs (including the 11 core ASVs) found in *Melipona* bee populations. Bootstrap values are shown in blue letters. The 11 core ASVs are written in bold characters. ^T^ Type strain. Trees are shown for the most abundant and core ASVs of A) *Apilactobacillus*, B) *Lactobacillus*, C) Streptococcaceae, D) Bifidobacteriaceae, and E) Acetobacteraceae. The phylogenetic trees were rooted according to the outgroups: (A) *Fructilactobacillus fructivorans*, (B) *Amylolactobacillus amylophilus,* (C) *Lactiplantibacillus plantarum*, (D) *Bombiscardovia coagulans*, (E) *Granulibacter bethesdensis.*

